# Set to its place: first DNA data on freshwater tardigrades of Sakhalin (Far East Russia) shed light on the phylogenetic position of *Mixibius* (Eutardigrada: Hypsibioidea)

**DOI:** 10.64898/2025.11.28.691205

**Authors:** Denis V. Tumanov

## Abstract

In this paper I present the results of the first faunistic investigation of the Sakhalin freshwater tardigrade fauna with the use of the integrative taxonomy. For the analysis, samples of bottom sediments were collected in several rivers of South Sakhalin. I found eight species belonging to three superfamilies of Eutardigrada. For most of those species I obtained data on mitochondrial COI gene and on 18S rRNA, 28 rRNA and ITS-2 sequences in addition to the morphological analysis using light and scanning electron microscopy. The number of tardigrade species known for Sakhalin Island increased from 1 to 9. Among 8 species found during my research one is similar to *Dactylobiotus dervizi* previously known from Komandorskiye Islands and one to *Murrayon hastatus*. Other 6 species belonging to the genera *Mixibius*, *Hypsibius*, *Dianea* and *Mesobiotus* are new for science, yet their formal description is not possible because of the small number of specimens. Inclusion of obtained molecular data in the phylogenetic analysis of the superfamily Hypsibioidea revealed for the first time the presence of the well-supported clade of the enigmatic genus *Mixibius* and gave possibility to recognise its phylogenetic position as sister clade to the genus *Acutuncus* within the family Acutuncidae. Two species of the genus *Mixibius* (*M. parvus* and *M. tibetanus*) are transferred into the genus *Isohypsibius* on the basis of the morphological analysis. Amended diagnoses for the family Acutuncidae and the genus *Acutuncus* are given.

## Introduction

Tardigrades are a group of microscopic segmented animals widely distributed in nature (Nelson *et al*. 2018). More than 1500 tardigrade species are known today (Degma & Guidetti 2025). The bulk of their species diversity is associated with semiterrestrial habitats containing thin-film of water (moss, lichens, soil etc.), however, they inhabit marine and freshwater biotopes as well (Nelson *et al*. 2018). Aquatic tardigrade species usually lack the ability to enter anhydrobiosis which is a well-known feature of semiterrestrial tardigrades (Nelson *et al*. 2018). This peculiarity of their biology hampered the use of the molecular taxonomic methods, especially in case of remote areas. Fixation of the total volume of the bottom substrates cannot provide sufficient extraction of the DNA from the obtained specimens. The best possible way to collect the material suitable for further studies is to extract individual specimens from the fresh samples directly in the field and fix them there. In the case of microscopic animals like tardigrades, such a method requires the field station equipped with optical and laboratory equipment. An alternative way is to freeze unprocessed samples and transport them to the laboratory. Both options are rarely possible during expeditions. Thus, the freshwater tardigrade fauna is much better investigated in those regions and locations that can be easily accessed from the stationary laboratories, mainly in Europe and North America. The only published DNA barcode record for the Asian freshwater tardigrade is for *Dactylobiotus taiwanensis* Camarda, Pai, Kristensen & Stec, 2025 so far.

Tardigrade fauna of the Russian Far East is still poorly investigated. There are only three publications devoted to the freshwater tardigrades of this region.

The first one is the paper by Ramazzotti (1966) with a brief description of the material from Lake Baikal. In this work he reported two unidentified species from the genera *Hypsibius* Ehrenberg, 1848 and *Macrobiotus* Schultze, 1834 (based on his brief text description, the latter species should now be attributed to the genus *Paramacrobiotus* Guidetti, Schill, Bertolani, Dandekar & Wolf, 2009). He also recorded *Thulinius augusti* (Murray, 1907b) (as *Hypsibius* (*Isohypsibius*) *augusti*), but after the revision of the genus *Thulinius* Bertolani, 2003, all previous reports of this species should be considered questionable (Bertolani *et al*. 1999). This work also included the description of *Grevenius baicalensis* (Ramazzotti, 1966) (as *Hypsibius* (*Isohypsibius*) *baicalensis*).

The second is the paper by Biserov (1992) with the description of two species from Baikal Lake: *Bertolanius markevichi* (Biserov, 1992) (as *Amphibolus* Bertolani, 1981) and *Vladimirobius irregibilis* (Biserov, 1992) (as *Isohypsibius* Thulin, 1928).

The last contribution is a brief conference abstract (Vvedenskaya 2009) devoted to the fauna of freshwater tardigrades of Kamchatka. She listed 14 species, but without any additional data it is impossible to evaluate the accuracy of the species identification.

All of these studies were published prior to the significant changes made by the introduction of molecular methods to the taxonomy of tardigrades. Currently all the records of the old “widely distributed” species should be considered questionable until confirmed using DNA barcoding methods (Gąsiorek 2024), which means that the investigation of tardigrade fauna should be started *de novo*, using molecular methods.

Sakhalin is the largest island in Russia, located in the Far East region. Tardigrade fauna of the island remains almost completely unknown. The only publication devoted to the Sakhalin tardigrades is the paper of Abe (2004) in which he described a new species *Parahypsibius stiliferus* (Abe, 2004) (as *Hypsibius*) from several locations all over the island.

During August 2024 I participated in a complex biological expedition on the Cape Crillon (South Sakhalin) organized by the non-profit charitable foundation “Support of bioresearch “BIOM”. During the field work numerous samples of the bottom sediments were collected from the rivers around the field station on Cape Anastasia. A few of them contained tardigrades and their eggs.

This paper is the first contribution to the knowledge of tardigrades inhabiting fresh waters of Sakhalin and the first investigation of the Sakhalin tardigrade fauna using DNA barcoding.

## Material and methods

### Sampling

Samples of the bottom sediments (sand, leaf litter, debris) were collected manually from the downstream of three rivers (Anastasia, Vodopadnaya and Atlasovka) and from small unnamed streams in the vicinity at these rivers’ mouth. Only samples collected from Vodopadnaya river and from a small stream near Atlasovka river contained tardigrades (Table 1). Upon collection, samples were transferred to the field laboratory where the tardigrade specimens and eggs were extracted using the standard technique of washing the sediment through two sieves with ≈ 1 mm mesh size and then with 29 μm mesh size (Tumanov 2018). The contents of the finer sieve were examined under a MBS-10 stereomicroscope, and obtained tardigrade specimens and eggs were picked up using a glass pipette. Considering the small number of specimens found (see Table 1), all material was fixed in RNA*later*™ Stabilization Solution (Thermo Fisher Scientific, USA), stored in a freezer and later transported to the laboratory at Saint-Petersburg State University.

**Table 1.** Sampling sites data. “V” indicates that the voucher specimen was used for LM after DNA extraction/.

| Location number | Location | Species | Number of specimens |  |  |
| --- | --- | --- | --- | --- | --- |
|  |  |  | LM | SEM | Genotyping |
| 1 | 46°0'20.16", 142°7'51.96", Cape Crillon, Cape Anastasia, small stream near Atlasovka river, below the waterfall, silted sand near the shore, coll. D.V. Tumanov, 16.08.2024 | *Hypsibius* sp. 1 | 1 (V) | – | 1 |
| 2 | 46°0'51.84", 142°9'38.16" Cape Crillon, Cape Anastasia, Vodopadnaya river, stream below the waterfall, silted sand near the shore, coll. D.V. Tumanov, 17.08.2024 | *Mesobiotus* sp. | 1 (V) | – | 1 |
|  |  | *Hypsibius* sp. 2 | 1 (V) | – | 1 |
|  |  | *Hypsibius* sp. 3 | 1 (V) | – | 1 |
| 3 | 46°0'52.20", 142°9'38.16" Cape Crillon, Cape Anastasia, Vodopadnaya river, small dam over the waterfall, silted sand near the shore, coll. D.V. Tumanov, 18.08.2024 | *Dianeia* sp. | 1 (V) | – | 1 |
|  |  | *Hypsibius* sp. 2 | 1 (V) | – | 1 |
|  |  | *Mixibius* sp. | 1 (V) | – | 1 |
| 4 | 46°0'51.84", 142°9'38.16" Cape Crillon, Cape Anastasia, Vodopadnaya river, waterfall, moss on the rock in the water stream, coll. D.V. Tumanov, 18.08.2024 | *Mesobiotus* sp. | 13 eggs | 5 eggs | 3 eggs |
| 5 | 46°0'51.84", 142°9'38.16" Cape Crillon, Cape Anastasia, Vodopadnaya river, stream below the waterfall, silted sand near the shore, coll. D.V. Tumanov, 19.08.2024 | *Dactylobiotus* cf. *dervizi* | 5 eggs | – | hatchlings from 3 eggs |
|  |  | *Murrayon* cf. *hastatus* | – | 5 eggs | – |

Photographs of the type material of *Dactylobiotus dervizi* Biserov, 1998 were used as the comparative material.

### Genotyping

DNA was extracted from individual tardigrade specimens and eggs using QuickExtract^™^ DNA Extraction Solution (Lucigen Corporation, USA, see complete protocol description in Tumanov (2020). Four markers were sequenced: a fragment of the cytochrome oxidase subunit I (COI) gene, the internal transcribed spacer (ITS-2), and fragments of the small (18S rRNA) and large (28S rRNA) ribosomal subunit gene. PCR reactions included 5 μl template DNA, 1 μl of each primer, 1 μl dNTP, 5 μl Taq Buffer (10X) (−Mg), 4 μl 25 mM MgCl_2_ and 0.2 μl Taq DNA Polymerase (Thermo Scientific^™^) in a final volume of 50 μl. The primers and PCR programs used are provided in Table 2. The PCR products were visualized in 1.5% agarose gel stained with ethidium bromide. All amplicons were sequenced directly using the ABI PRISM Big Dye Terminator Cycle Sequencing Kit (Applied Biosystems) with the help of an ABI Prism 310 Genetic Analyzer in the Core Facilities Center “Centre for Molecular and Cell Technologies” of St. Petersburg State University. Sequences were edited and assembled using ChromasPro software (Technelysium). The COI sequences were translated to amino acids using the invertebrate mitochondrial code, implemented in MEGA11 (Tamura *et al*. 2021), in order to check them for the presence of stop codons and therefore of pseudogenes. A complete list of the obtained gene sequences is presented in Table 3.

**Table 2.**
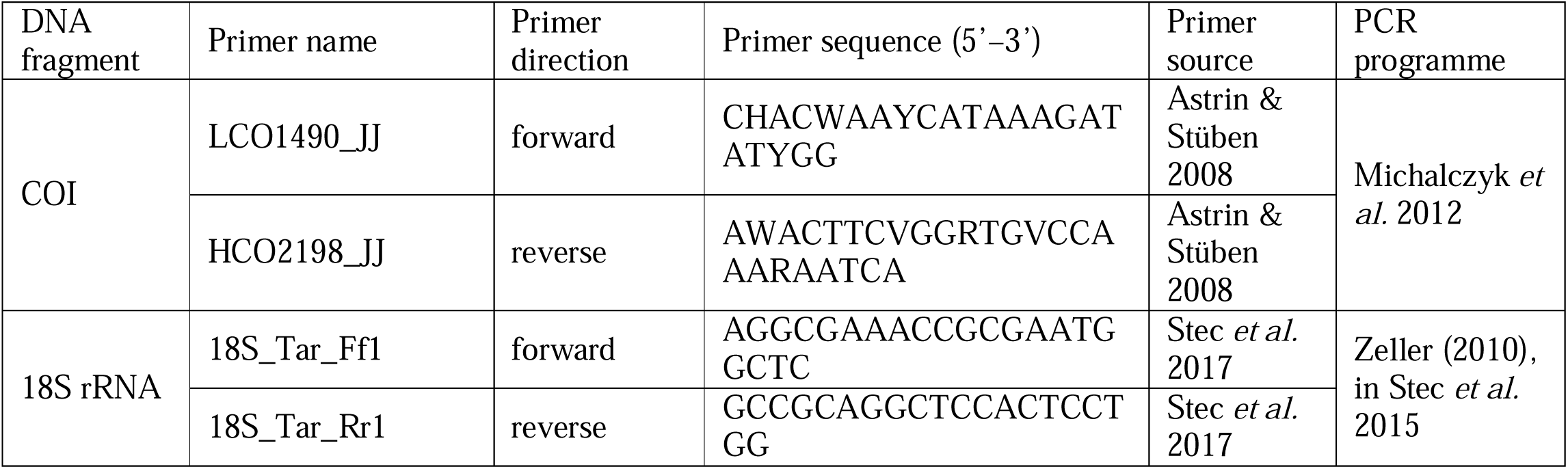
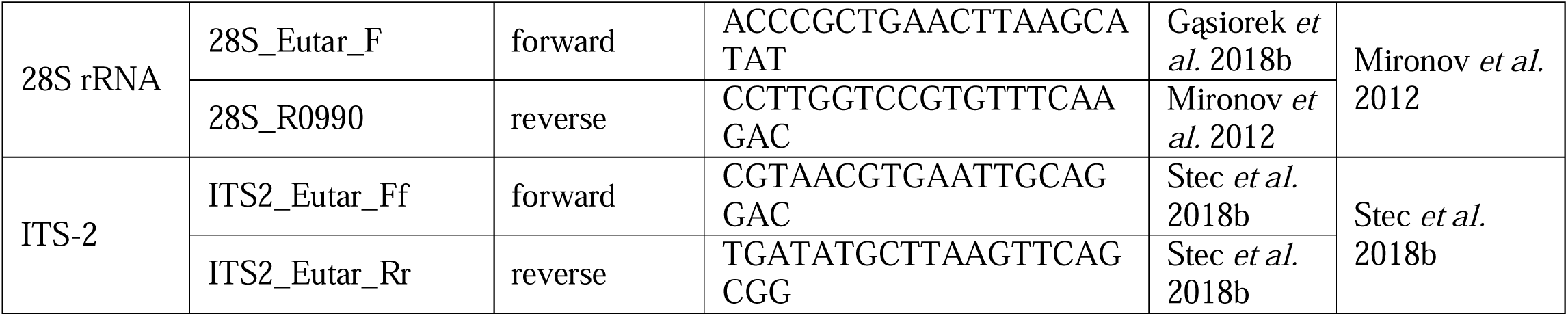
Primers and PCR programs used for amplification of the four DNA fragments sequenced in the study.

**Table 3.** List of gene sequences obtained in the study.

| Species | Specimen Catalog # | Genes |  |  |  |
| --- | --- | --- | --- | --- | --- |
|  |  | 18S rRNA | 28S rRNA | ITS-2 | COI |
| *Mixibius* sp. | SPbSU 370(003) | PX243403 | PX232532 | NA | PX233160 |
| *Hypsibius* sp. 1 | SPbSU 369(001) | PX243404 | PX232533 | PX232521 | PX233161 |
| *Hypsibius* sp. 2 | SPbSU 370(002) | PX243405 | PX232534 | PX232522 | PX233162 |
| *Hypsibius* sp. 2 | SPbSU 374(002) | NA | PX232535 | PX232523 | PX233163 |
| *Hypsibius* sp. 3 | SPbSU 374(003) | PX243406 | PX232536 | PX232524 | PX233164 |
| *Dianeia* sp. | SPbSU 370(001) | NA | PX232537 | NA | PX233165 |
| *Mesobiotus* sp. | SPbSU 374(001) | NA | PX232541 | PX232528 | PX233169 |
| *Mesobiotus* sp. | SPbSU 375(001) | PX243410 | PX232542 | PX232529 | PX233170 |
| *Mesobiotus* sp. | SPbSU 375(002) | PX243411 | PX232543 | PX232530 | PX233171 |
| *Mesobiotus* sp. | SPbSU 375(003) | PX260307 | NA | PX232531 | PX233172 |
| *Dactylobiotus* cf. *dervizi* | SPbSU 366(001, 002) | PX243407 | PX232538 | PX232525 | PX233166 |
| *Dactylobiotus* cf. *dervizi* | SPbSU 367(001, 002) | PX243408 | PX232539 | PX232526 | PX233167 |
| *Dactylobiotus* cf. *dervizi* | SPbSU 367(004) | PX243409 | PX232540 | PX232527 | PX233168 |

### Microscopy and imaging

Following the DNA extraction, tardigrade cuticles and eggshells were mounted on slides in Hoyer’s medium. Permanent slides were examined under a Leica DM2500 microscope equipped with phase contrast (PhC) and differential interference contrast (DIC). Photographs were taken using a Nikon DS-Fi3 digital camera with NIS software. In the case of deep structures that could not be fully focused in a single image, a series of several photographs were taken and then stacked into a single deep-focus image using Helicon Focus software (Helicon Soft Ltd).

For SEM investigation, eggs were dehydrated in an ascending ethyl alcohol series (10%, 20%, 30%, 50%, 70%, 96%) and acetone, critical-point dried in CO_2_, mounted on stubs, and coated with gold. A Tescan MIRA3 LMU scanning electron microscope (Tescan, Brno, Czech Republic) and Hitachi TM-1000 (Hitachi, Japan) were used for observations.

### Phylogenetic analyses

In addition to the newly obtained sequences, we used sequences of 18S and 28S rRNA genes available in GenBank for the Hypsibioidea species (accessed on 5 July 2025), which (i) had appropriate length (more than 600 bp for 18S rRNA, all available 28S rRNA sequences), (ii) were homologous to the newly obtained sequences and (iii) originated from publications with a reliable attribution of the investigated taxa (see SM.01). Both 18S rRNA and 28S rRNA are nuclear markers used in phylogenetic analyses to distinguish higher taxonomic levels (Guil & Giribet 2012; Bertolani *et al*. 2014; Guil *et al*. 2019; Gąsiorek *et al*. 2019, 2023; Gąsiorek & Michalczyk 2020; Tumanov 2022; Tumanov & Tsvetkova 2023; Vecchi *et al*. 2023). The COI and ITS-2 markers were not included in the analysis due to the negative effect on the deep phylogeny resolution that may give the inclusion of fast-evolving genes (Betancur-R *et al*. 2014; Chen *et al*. 2015; Klopfstein *et al*. 2017; Tumanov 2022; Tumanov & Tsvetkova 2023). Two sequences of Macrobiotoidea (*Macrobiotus shonaicus* Stec, Arakawa & Michalczyk, 2018a, Macrobiotidae and *Richtersius coronifer* (Richters, 1903), Richtersiusidae) were used as an outgroup.

Sequences were automatically aligned with the MAFFT algorithm (Katoh *et al*. 2002) using AliView software version 1.27 (Larsson 2014); alignments were manually trimmed to lengths of 1677 bp for 18S and 826 bp for 28S in order to exclude uncertainty in the terminal regions of sequences. The sequences of both genes were concatenated using SeaView 4.0 (Gouy *et al*. 2010) (see final alignment in SM.02). The best substitution model and partitioning scheme for posterior phylogenetic analysis were selected under the Akaike information criterion using IQ-TREE multicore version 1.6.12 (Kalyaanamoorthy *et al*. 2017; Minh *et al*. 2020). IQ-TREE suggested retaining two predefined partitions separately; the SYM+I+G4 model was found to be the best fit for the first dataset (18S rRNA) and the GTR+F+I+G4 model was found to be the best fit for the second dataset (28S rRNA). Maximum-likelihood (ML) topologies were constructed using IQ-TREE software (Minh *et al*. 2020). Bayesian analysis of the same datasets was performed using MrBayes version 3.2.6, a GTR model with gamma correction for intersite rate variation (eight categories) and the covariation model (Ronquist & Huelsenbeck 2003). I ran the analyses as four separate chains (default heating parameters) for 1000000 generations, by which time they had ceased to converge (the final average SD of the split frequencies was less than 0.01). The quality of chains was estimated using built-in MrBayes tools. The quality of Bayesian analysis was verified using the program Tracer v1.7.1 (Rambaut *et al*. 2018). Unadjusted pairwise distances were calculated using MEGA11 (Tamura *et al*. 2021) with the treatment of gaps/missing data set to “pairwise deletion”. The homology comparison of the obtained sequences with GenBank records was performed using BLASTn algorithm (https://blast.ncbi.nlm.nih.gov/Blast.cgi). All final Bayesian and ML consensus trees were visualised using FigTree v.1.4.4 (Rambaut 2018). Species delimitation was done on the COI alignments for the genera *Hypsibius* (including *Borealibius* and *Cryobiotus*) and *Mesobiotus*, and the ITS-2 alignment for the genus *Hypsibius* (including *Borealibius* and *Cryobiotus*) using bPTP web server https://species.h-its.org/ (Zhang *et al*. 2013).

### Institutional acronyms

Specimens from the following institutions and collections were examined (the curator is given in parenthesis).

SPbSU = St. Petersburg State University, Russia, Faculty of Biology, Department of Invertebrate Zoology (Denis Tumanov).

## Results

Phylum Tardigrada Doyère, 1840

Class Eutardigrada Richters, 1926

Order Parachela Schuster, Nelson, Grigarick & Christenberry, 1980

### Superfamily Hypsibioidea Pilato, 1969

#### Incerta familia

##### Genus *Mixibius* Pilato, 1992

***Mixibius* sp. (Fig. 1).** Location 3, one adult specimen. Morphology of the adult specimen mainly conforms to the diagnoses of both *Acutuncus* Pilato & Binda, 1997 and *Mixibius*: pharynx with two macroplacoids, without microplacoids or septulum, external claws of the *Isohypsibius* type, internal claw of the *Hypsibius* type. Morphology of the apophyses for the insertion of the stylet muscles (AISM) could not be categorized because of the buccal tube position on the slide.

**FIGURE 1.**
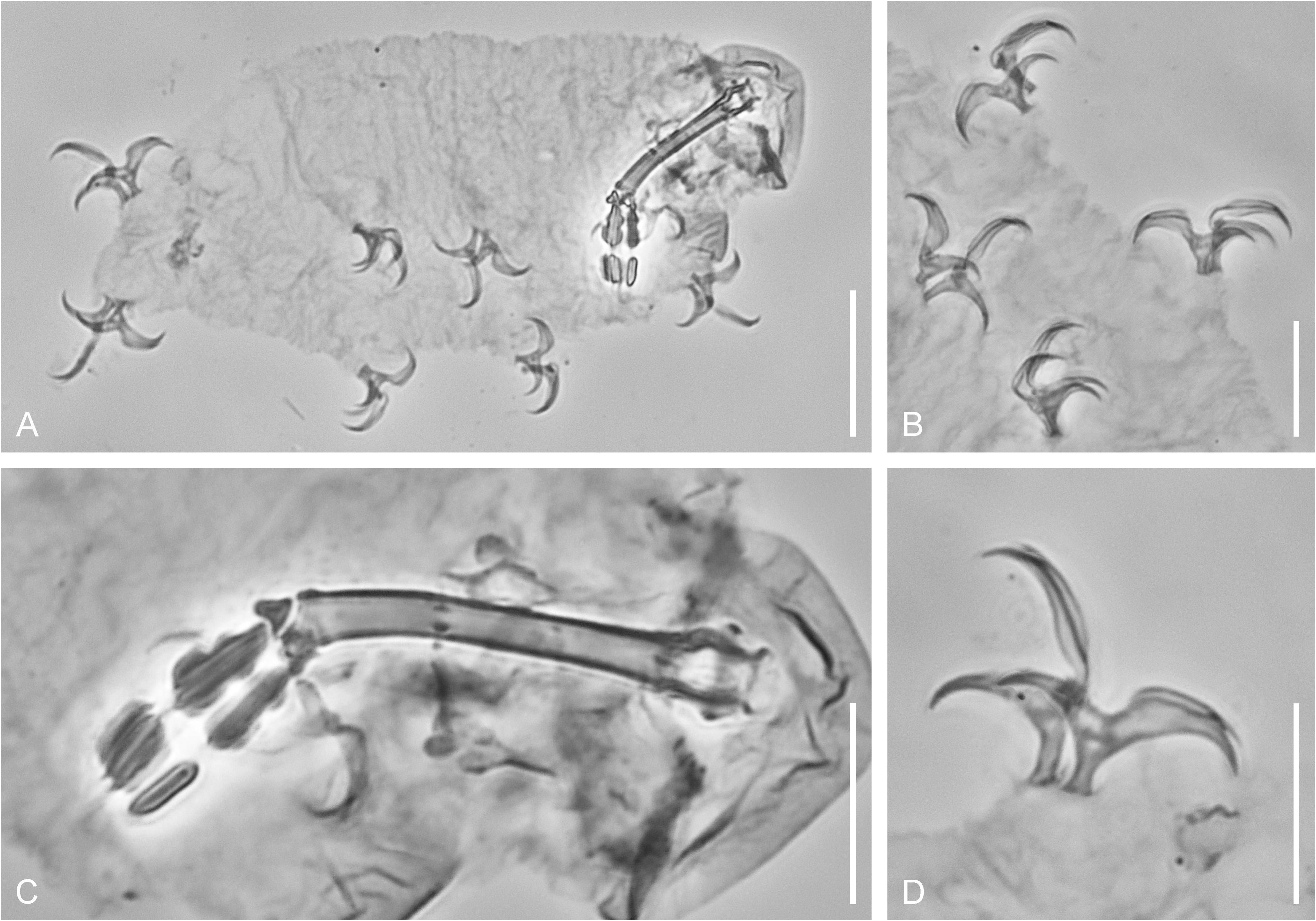
*Mixibius* sp., details of morphology in PhC. A. Voucher specimen after DNA extraction (lateral view). B. Claws of legs I–II. C. Buccal-pharyngeal apparatus. D. Claws of leg IV. Scale bars: A = 20 µm, B–D = 10 µm.

Sequences of three genes were obtained for the specimen: 18S rRNA, 28S rRNA, and COI (see Table 3), an attempt to amplify the ITS-2 marker was unsuccessful. Homology comparison of the obtained sequences with GenBank records (23^rd^ November 2025) revealed affinity to the superfamily Hypsibioidea. BLASTn percent identity for 18S and 28S sequences of the studied specimen to the sequences of *Acutuncus antarcticus* (Richters 1904) was 98.8% (EU266944, Sands *et al*. 2008) and 93.2% (OM278643, Vecchi *et al*. 2023) respectively.

### Family Hypsibiidae Pilato, 1969

#### Subfamily Hypsibiinae Pilato, 1969

##### Genus *Hypsibius* Ehrenberg, 1848

***Hypsibius* sp. 1 (Fig. 2).** Location 1, one adult specimen. Morphology of the adult specimen conforms to the diagnosis of the *Hypsibius dujardini* (Doyère, 1840) morphogroup, recognized here as a group of *Hypsibius* species with a smooth or slightly rugose (but not granulate or tuberculate) cuticle, two macroplacoids and a septulum in the pharynx (Tumanov & Avdeeva 2021). Sequences of four markers were obtained for the specimen: 18S rRNA, 28S rRNA, ITS-2, and COI (see Table 3). Homology comparison of the obtained sequences with GenBank records (5^th^ July 2025) indicated similarity to the genus *Hypsibius*. The range of uncorrected genetic *p*-distances for COI and ITS-2 sequences between the studied specimen and sequences of the genus *Hypsibius* (with addition of the closely related genera *Borealibius* Pilato, Guidetti, Rebecchi, Lisi, Hansen & Bertolani, 2006 and *Cryobiotus* Dastych, 2019) available in GenBank are presented in SM.03. Considering the significant genetic distance for the COI sequence from all known *Hypsibius* species (the closest is *Hypsibius* cf. *exemplaris* from Svalbard; MW010373; Zawierucha *et al*. 2020, *p*-distance is 14.8%) this species seems to be a potentially cryptic species in the *Hypsibius dujardini* morphogroup. This conclusion was supported by the results of the bPTP species delimitation (see SM.04).

**FIGURE 2.**
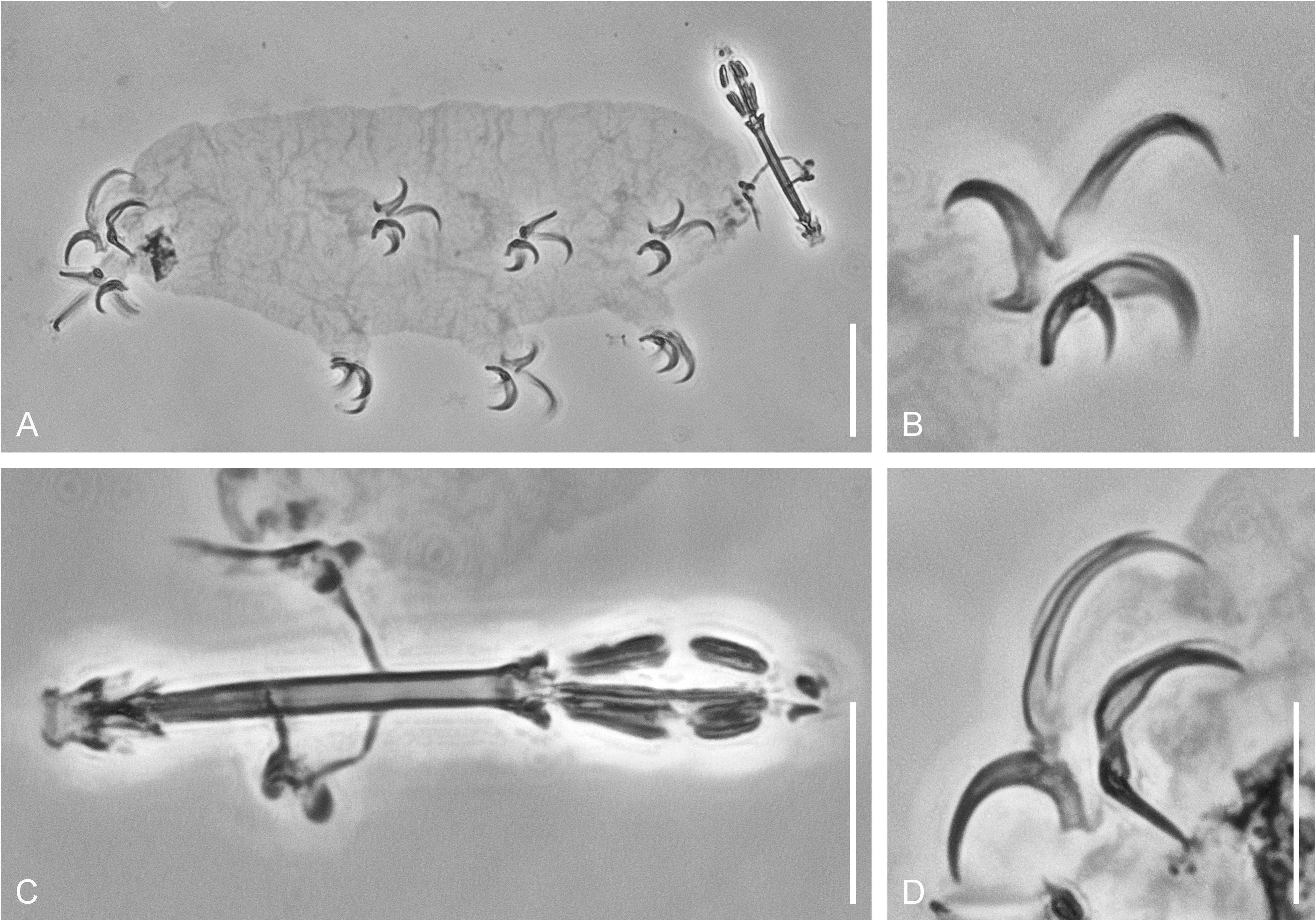
*Hypsibius* sp. 1, details of morphology in PhC. A. Voucher specimen after DNA extraction (lateral view). B. Claws of leg II. C. Buccal-pharyngeal apparatus. D. Claws of leg IV. Scale bars: A = 20 µm, B–D = 10 µm.

***Hypsibius* sp. 2 (Fig. 3).** Locations 2 and 3, two adult specimens. Morphology of both specimens conforms to the diagnosis of the *Hypsibius dujardini* morphogroup. Sequences of four markers were obtained: 18S rRNA, 28S rRNA, ITS-2, and COI (see Table 3). Presence of two haplotypes of the COI gene was revealed, with 0.99% *p*-distance between them. Homology comparison of the obtained sequences with GenBank records (5^th^ July 2025) indicated similarity to the genus *Hypsibius*. The range of uncorrected genetic *p*-distances for COI and ITS-2 sequences between the studied specimen and sequences of the genus *Hypsibius* (with addition of the closely related genera *Borealibius* and *Cryobiotus*) available in GenBank are presented in SM.03. Considering the significant genetic distance for the COI sequence from all known *Hypsibius* species (the most close is *Hypsibius* sp. from Italy; PQ140627; Surmacz *et al*. 2025, *p*-distances are 13.6% and 12.8% for each haplotype respectively) and from *Hypsibius* sp. 1 described above (16.4% and 18.0% for each haplotype respectively) this species seems to be another potentially cryptic species in the *Hypsibius dujardini* morphogroup found in Sakhalin. This conclusion was supported by the results of the bPTP species delimitation (see SM.04).

**FIGURE 3.**
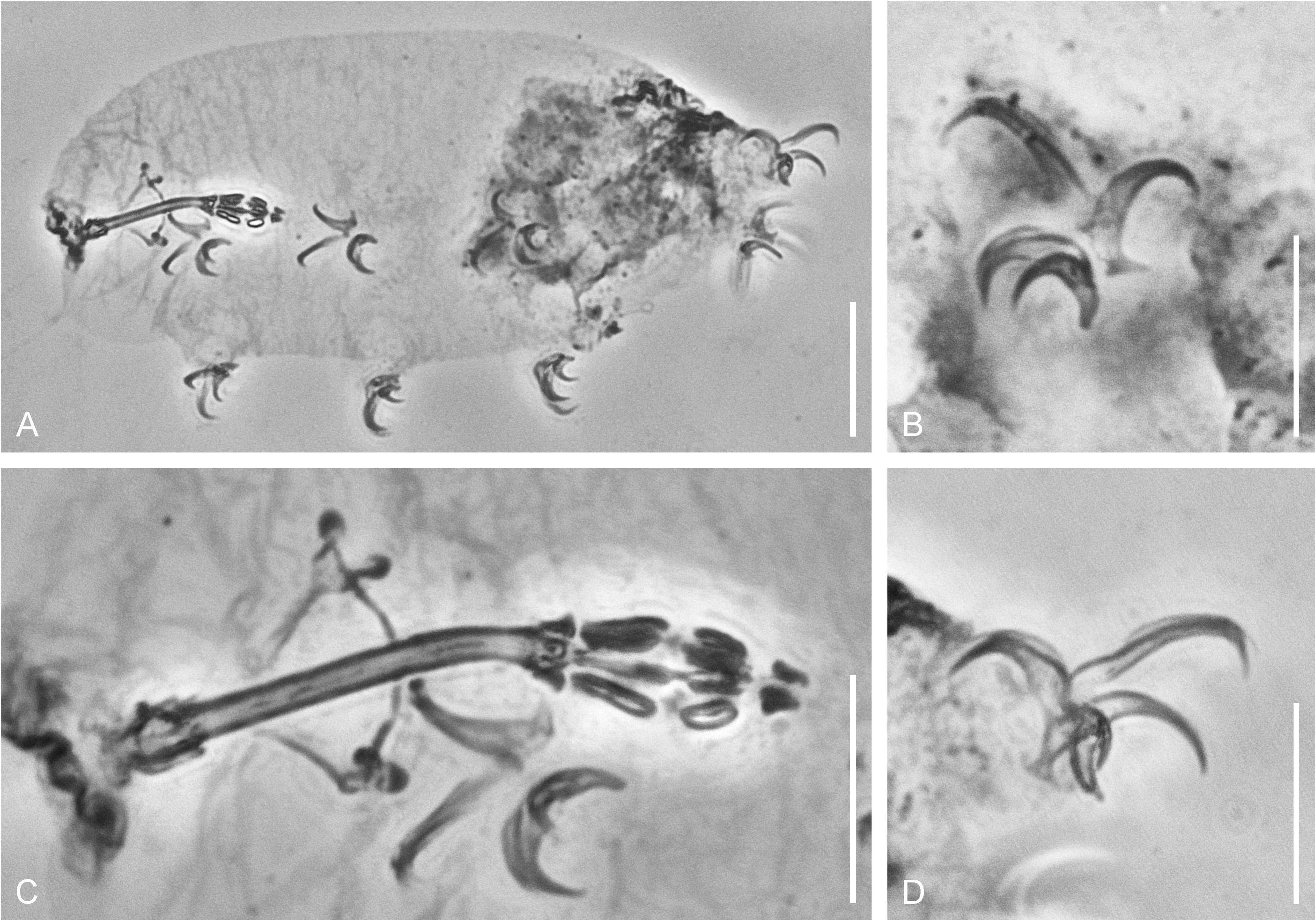
*Hypsibius* sp. 2, details of morphology in PhC. A. Voucher specimen after DNA extraction (lateral view). B. Claws of leg III. C. Buccal-pharyngeal apparatus. D. Claws of leg IV. Scale bars: A = 20 µm, B–D = 10 µm.

***Hypsibius* sp. 3 (Fig. 4).** Location 2, one adult specimen. Morphology of the adult specimen conforms to the diagnosis of the *Hypsibius convergens* (Urbanowicz, 1925) morphogroup, recognized here as a group of *Hypsibius* species with a smooth or slightly rugose (but not granulate or tuberculate) cuticle, two elongate macroplacoids without septulum (or microplacoid) in the pharynx (Gąsiorek *et al*. 2024). Sequences of four markers were obtained for the specimen: 18S rRNA, 28S rRNA, ITS-2, and COI (see Table 3). Homology comparison of the obtained sequences with GenBank records (5^th^ July 2025) indicated similarity to the genus *Hypsibius*. The range of uncorrected genetic *p*-distances for COI and ITS-2 sequences between the studied specimen and sequences of the genus *Hypsibius* (with addition of the closely related genera *Borealibius* and *Cryobiotus*) available in GenBank are presented in SM.03. Considering the significant genetic distance for the COI sequence from all known *Hypsibius* species (the closest is *Hypsibius* sp. from Poland; PQ140631–2; Surmacz *et al*. 2025, *p*-distance is 17.2%) this species is the third new species of the genus *Hypsibius* found in Sakhalin. This conclusion was supported by the results of the bPTP species delimitation (see SM.04).

**FIGURE 4.**
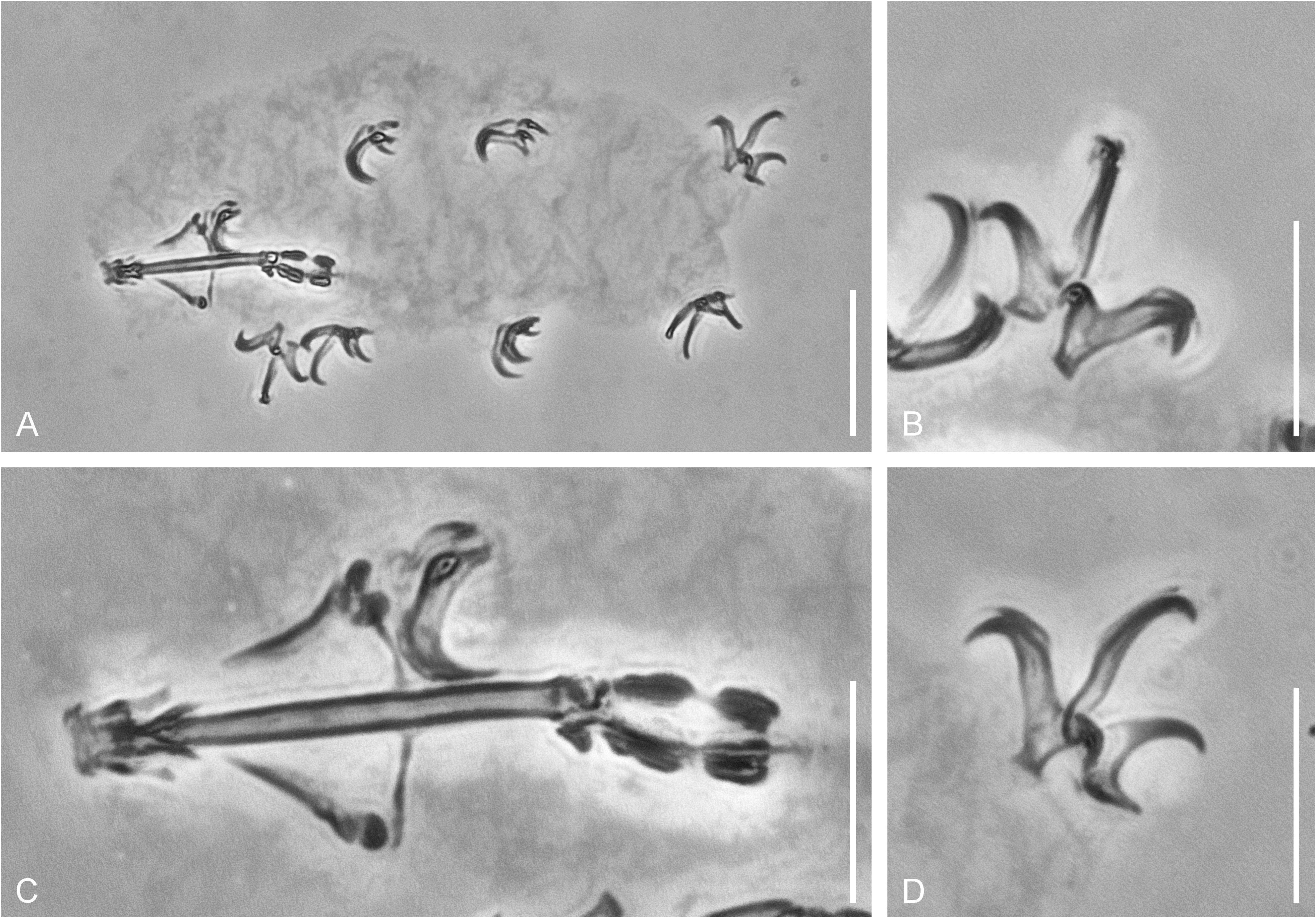
*Hypsibius* sp. 3, details of morphology in PhC. A. Voucher specimen after DNA extraction (dorso-ventral view). B. Claws of leg I. C. Buccal-pharyngeal apparatus. D. Claws of leg IV. Scale bars: A = 20 µm, B–D = 10 µm.

### Superfamily Isohypsibioidea Sands *et al*., 2008

#### Family Isohypsibiidae Sands et al., 2008

#### Genus *Dianea* Gąsiorek, Stec, Morek & Michalczyk, 2019

***Dianea* sp. (Fig. 5).** Location 3, one adult specimen. Morphology of the adult specimen conforms to the diagnosis of the genus *Dianea* (Gąsiorek *et al*. 2019) and the redescription of *Dianea sattleri* (Richters, 1902) (Dastych 1990). Sequences of two markers were obtained for the specimen: 28S rRNA and COI (see Table 3), attempts to amplify 18S rRNA and ITS-2 markers were unsuccessful. Homology comparison of the obtained sequences with GenBank records (5^th^ July 2025) indicated similarity to the genus *Dianea*. Uncorrected *p*-distance for COI sequences between the studied specimen and the single sequence available in GenBank (*Dianea* sp. from Poland; PQ140621; Surmacz *et al*. 2025) is 17.2%. Considering this significant genetic distance, this specimen seems to represent a potentially cryptic new species of the genus *Dianea*.

**FIGURE 5.**
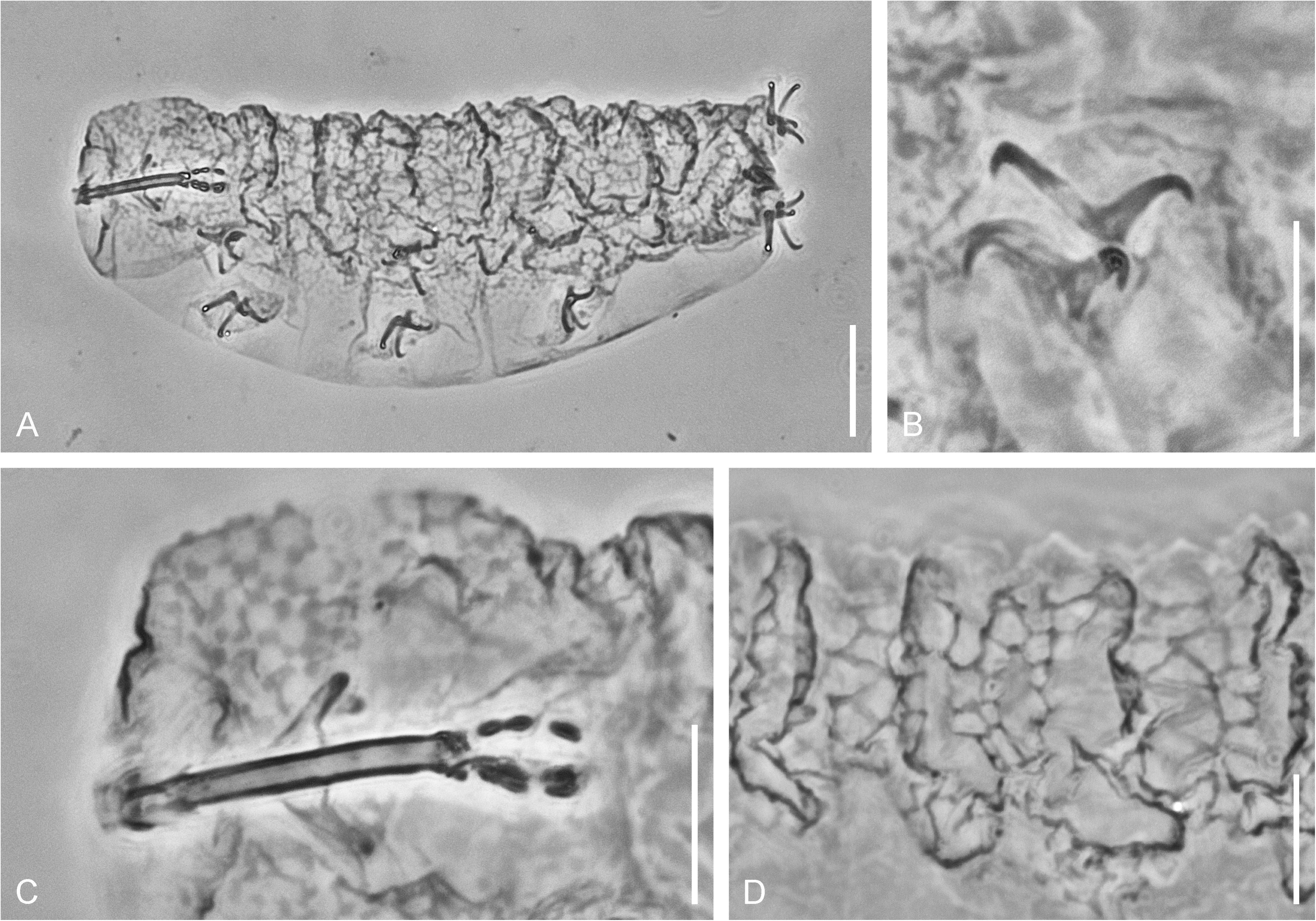
*Dianea* sp., details of morphology in PhC. A. Voucher specimen after DNA extraction (lateral view). B. Claws of leg III. C. Buccal-pharyngeal apparatus and cuticular sculpture of the cephalic region. D. Dorsal cuticular sculpture. Scale bars: A = 20 µm, B–D = 10 µm.

### Superfamily Macrobiotoidea Thulin, 1928

#### Family Macrobiotidae Thulin, 1928

##### Genus *Mesobiotus* Vecchi, Cesari, Bertolani, Jönsson, Rebecchi & Guidetti, 2016

***Mesobiotus* sp. (Fig. 6).** Location 2, one juvenile specimen; location 4, 13 eggs. Morphology of the specimen and the eggs are typical for the genus *Mesobiotus, M. harmsworthi* (Murray, 1907a) morphogroup (according Kaczmarek *et al*. 2020). Sequences of four markers were obtained for the single specimen and three eggs: 18S rRNA, 28S rRNA, ITS-2, and COI (see Table 3). Homology comparison of the obtained sequences with GenBank records (5^th^ July 2025) indicated similarity to the genus *Mesobiotus*. Despite the fact that the single specimen and the eggs were collected in different locations (which though belongs to the same waterbody), they were considered conspecific because of the similarity of the COI gene sequences (uncorrected *p*-distances between the juvenile specimen and the eggs is 2.4%, which is within the limits of intraspecific variability for tardigrades). The range of uncorrected genetic *p*-distances for COI and ITS-2 sequences between the studied specimens and sequences of the genus *Mesobiotus* available in GenBank are presented in SM.05. Considering the significant genetic distance for the COI sequences from all known *Mesobiotus* species (the closest is *Mesobiotus bockebodicus* Atherton, Hulterström, Guidetti & Jönsson, 2025 from Sweden; PQ365776; Atherton *et al*. 2025, *p*-distance is 19.8%), this is a new species of the genus *Mesobiotus*. This conclusion was supported by the results of the bPTP species delimitation (see SM.06).

**FIGURE 6.**
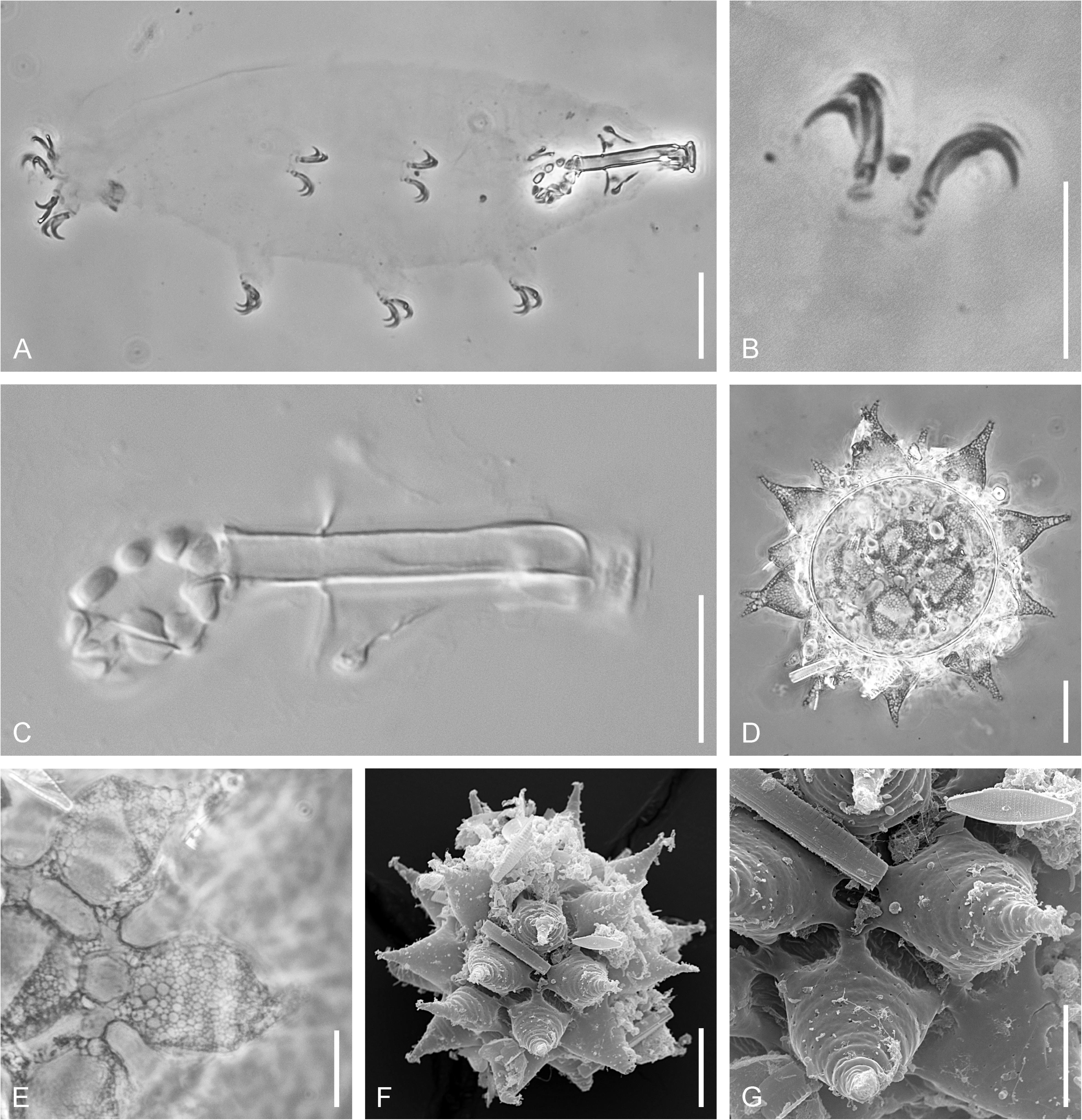
*Mesobiotus* sp., details of morphology. A. Voucher specimen after DNA extraction – in PhC (lateral view). B. Claws of leg II – in PhC. C. Buccal-pharyngeal apparatus – in DIC. D. Egg – in PhC. E. Details of the egg surface – in PhC. F. Egg – in SEM. G. Details of the egg surface – in SEM. Scale bars: A, D, F = 20 µm, B, C, G = 10 µm.

#### Family Murrayidae Guidetti, Rebecchi & Bertolani, 2000

##### Genus *Dactylobiotus* R.O. Schuster, 1980 (in Schuster et al. 1980)

***Dactylobiotus* cf*. dervizi* Biserov, 1998 (Fig. 7).** Location 5, five eggs. During processing of 3 eggs, initially fixed in RNA*later*™ in ddH_2_O, three juvenile specimens hatched from the eggs. All of them were processed to the extraction of DNA and then mounted on slides. Morphology of these specimens and eggs perfectly conforms to the description of *D. dervizi* from Komandorskiye Islands (Biserov 1998). Egg shell with numerous irregularly distributed small pores not only on the egg surface between processes, but also on the processes’ bases, forming wide band. This character was not mentioned in the original description, but the band of the pores is well visible in the original SEM photos (figures 13–14 in Biserov 1998) and in the reinvestigated type material eggs. There are two differences from the original description of *D. dervizi*: the presence of eyes in my material, while *D. dervizi* have no eyes according to Biserov (1998), and less frequent processes with branching apexes. Taking into account a small number of specimens analysed in the original description (5 adult animals), the absence of adult animals in my material, inconstancy of such character as presence or absence of eyes within Eutardigrada, and variability of egg shall characters in *Dactylobiotus* (Camarda *et al*. 2025), my material should be tentatively attributed as *Dactylobiotus* cf*. dervizi*.

**FIGURE 7.**
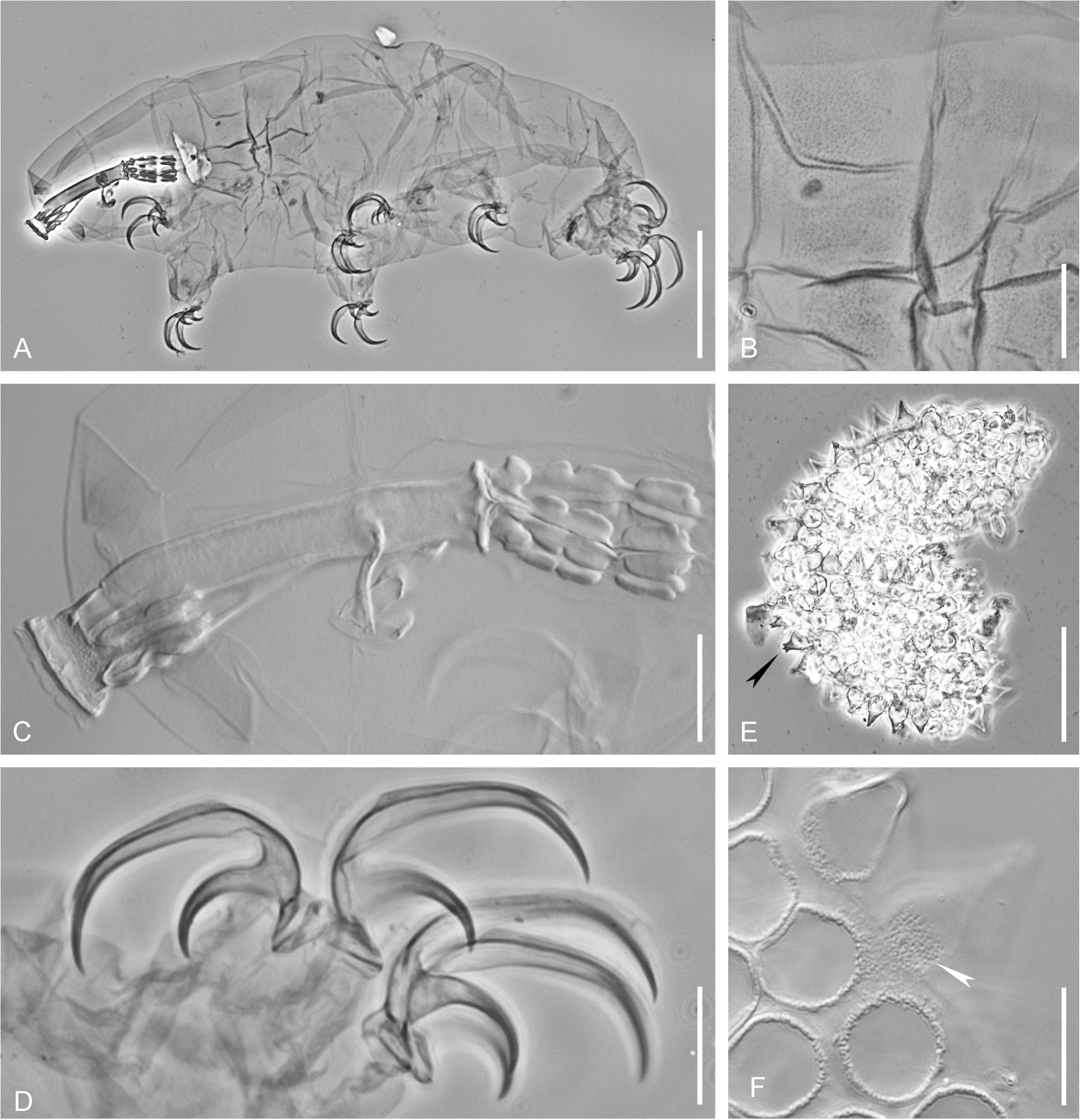
*Dactylobiotus* cf. *dervizi*, details of morphology. A. Voucher specimen after DNA extraction – in PhC (lateral view). B. Cuticular sculpture – in PhC. C. Buccal-pharyngeal apparatus – in DIC. D. Claws of leg IV – in PhC. E. Egg – in PhC, black arrowhead points to the branched process of the egg chorion. F. Details of the egg surface – in DIC, white arrowhead points to the pores in the basal part of the process of the egg chorion. Scale bars: A, E = 50 µm, B, C, D, F = 10 µm.

##### Genus *Murrayon* Bertolani & Pilato, 1988

***Murrayon* cf. *hastatus* (Murray, 1907) (Fig. 8).** Location 5, five eggs. The morphology of the eggs presented in the SEM images (Fig. 8A, B) is in concordance with the SEM observations of *M. hastatus* eggs collected in North-West Russia (Fig. 8C, D). Egg shell processes do not form continuous layer on the egg surface, but are arranged in meshes, consisted of 4–6 processes, connected with their lateral sides, with a deep pit in centre of each mesh. Eggs from Sakhalin differs from eggs of *M. hastatus* from North-West Russia in having no filamentous appendages on the top of each egg shell process, well developed in the eggs of *M. hastatus.* This can be an evidence for the presence of a new species of this group in the fauna of Sakhalin, but the presence or absence of such appendages is a variable character within the European populations of *M. hastatus.* Taking into account genetic variability between *M. hastatus* populations in European Russia (Tumanov, unpublished data), this morphospecies is likely a group of cryptic or pseudocryptic species waiting for a revision.

**FIGURE 8.**
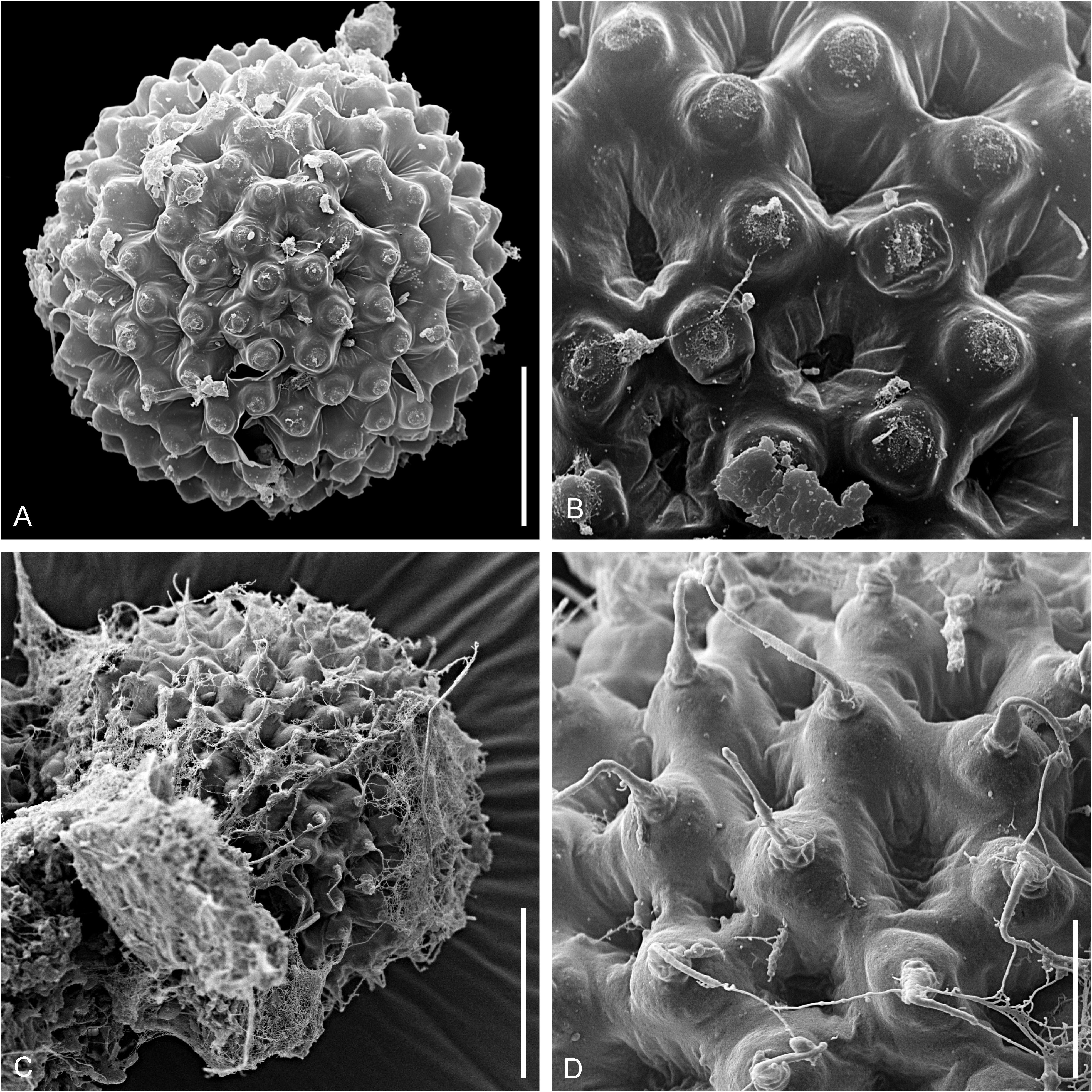
The egg of *Murrayon* cf. *hastatus* in comparison with *M*. *hastatus* in SEM. A. Total view of the egg of *M*. cf. *hastatus* from Sakhalin. B. Surface details of the same egg. C. Total view of the egg of *M*. *hastatus* from North-West Russia. D. Surface details of the same egg. Scale bars: A, B = 20 µm, C, D = 5 µm.

## Discussion

### Notes on the phylogenetic position of the genus *Mixibius*

Until now, representatives of the monotypic family Acutuncidae (comprising the single genus *Acutuncus*) were known only from the Western Palaearctic (Svalbard, Zawierucha *et al*. 2020; Italy and the United Kingdom, Vecchi *et al*. 2023) and Antarctica (Vecchi *et al*. 2023). The genus *Mixibius* shares the same type of claws with the genus *Acutuncus,* which represents a combination of *Hypsibius* and *Isohypsibius* types. Currently, *Mixibius* includes all species with this claw type, except those attributed to the genus *Acutuncus* on the basis of phylogenetic analysis and egg shell morphology (Vecchi *et al*. 2023). The type species of this genus, *Mixibius saracenus* (Pilato, 1973) was described from fresh-waters of Sicily (Europe) (Pilato 1973). Other species currently attributed to this genus are known from Europe (Biserov 1999; Lisi *et al*. 2014), Asia (Pilato *et al*. 2004, 2010; Li & Li 2008; Wang 2009), North America (Kaczmarek *et al*. 2016), South America (Kaczmarek *et al*. 2015), and Antarctica (Pilato *et al*. 2017). For the Far East of Russia, the genus *Mixibius* was recorded once, from several waterbodies of the Kamchatka Peninsula (Vvedenskaya 2009) as *M. saracenus*, but this identification needs verification. Only two sequences (each of the 18S rRNA gene) are available for this genus in GenBank – one for the type species and one attributed as *Mixibius* cf. *saracenus* (Bertolani *et al*. 2014). Both sequences were previously excluded from the phylogenetic analysis of the superfamily Hypsibioidea because of their unstable position in the resulting trees (Vecchi *et al*. 2023).

The phylogenetic analysis, which included both *Mixibius* sequences from GenBank and sequences obtained in the present work, revealed their close affinity to each other (Fig. 9). Monophyly of this clade was stable and strongly supported in Bayesian analysis (posterior probability value is 0.99), while poorly supported in ML (bootstrap value is 65). Within this clade, *Mixibius* sp. from Sakhalin is more closely related to the type species of the genus than to *Mixibius* cf. *saracenus* from Italy. This clade forms a sister group to the genus *Acutuncus* in both Bayesian and ML analyses, but also with weak support in the latter (0.99 and 68, respectively).

**FIGURE 9.**
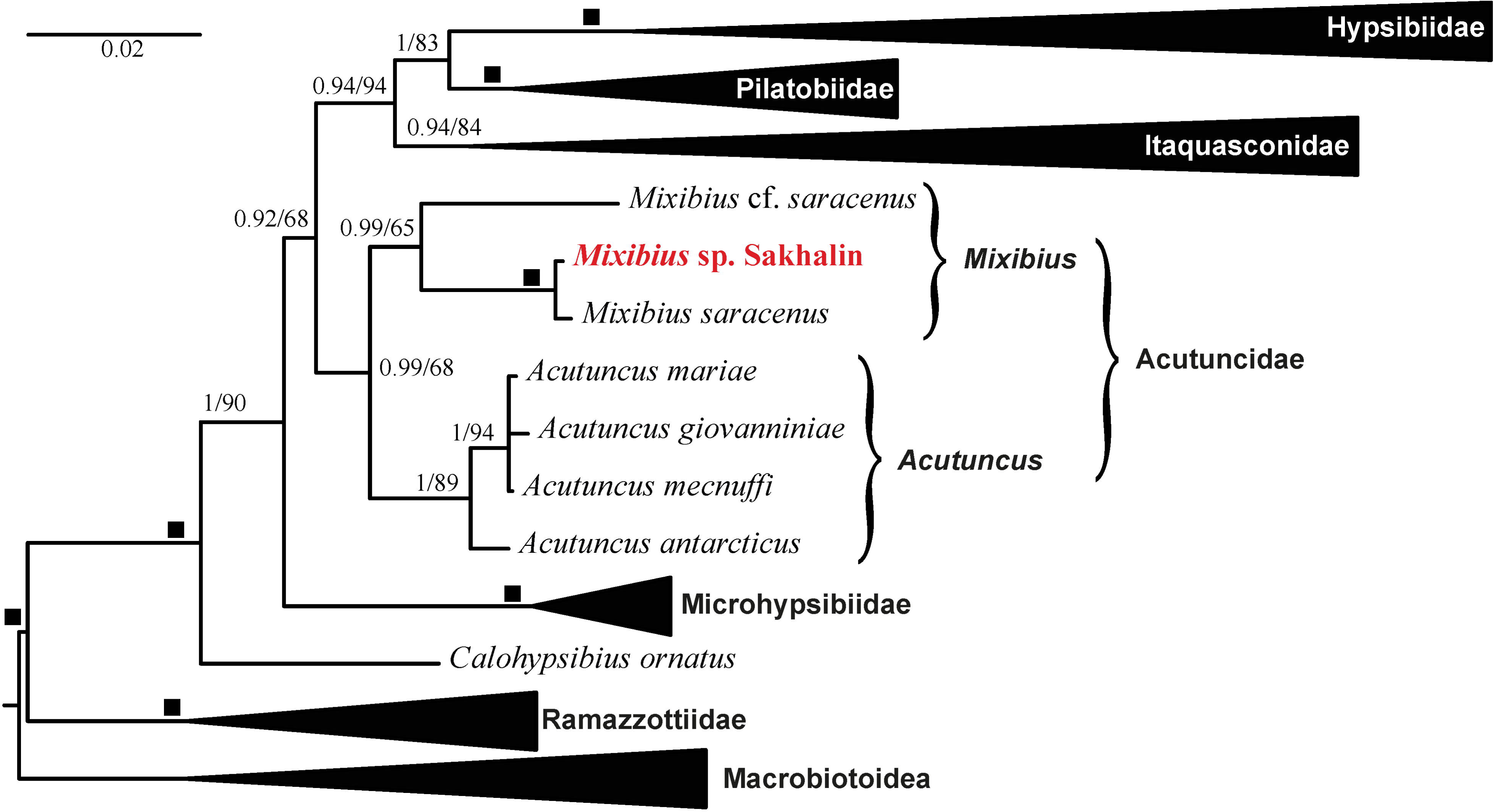
The phylogeny of Hypsibioidea based on concatenated 18S + 28S rRNA sequences. Numbers at nodes indicate Bayesian posterior probability values (BI, on the left) and bootstrap values (ML, on the right). Black squares indicate the nodes supported by values of 1/100% with both methods. Scale bar and branch lengths refer to the Bayesian analysis.

Until now, the genus *Mixibius* was considered *incerta familia* within Hypsibioidea (see Bertolani *et al*. 2014 and Vecchi *et al*. 2023). There are evident similarities between the genera *Mixibius* and *Acutuncus* in both morphological (the same type of buccal-pharyngeal apparatus and the same type of claws) and molecular characteristics (Fig. 9). In my opinion, the genus *Mixibius* should be included into the family Acutuncidae. Considering that it forms a clade distinct from the genus *Acutuncus*, it should be retained as an independent genus within the family Acutuncidae. All *Mixibius* species other than the type species (*M. saracenus*) and two unnamed species (*Mixibius* cf. *saracenus* from Italy and *Mixibius* sp. from Sakhalin) should be tentatively retained within the genus *Mixibius* until receiving new data. The exception is two species currently attributed to the genus *Mixibius* (*Mixibius parvus* Lisi, Sabella & Pilato, 2014 and *Mixibius tibetanus* Li & Li, 2008). Both have evidently different organisation of the buccal-pharyngeal apparatus with three macroplacoids and a microplacoid. Moreover, the shape of the claws of these species can not confirm their attribution to the genus *Mixibius*. The claws of *M. tibetanus* (see fig. 14 in Li & Li 2008) are of typical *Isohypsibius* type, indistinguishable from those of *Is. prosostomus* Thulin, 1928. The shape of the claws of *M. parvus* is not completely clear because of unfavorable position on the slide (see fig. 2 in Lisi *et al*. 2014). In addition, *M. tibetanus* possesses *Isohypsibius*-type cuticular bars near the claw bases on legs I–III and reticulate cuticular sculpture, which is widely distributed within the *I. prosostomus* complex. So, there are no trustworthy evidences supporting the inclusion of these species in the genus *Mixibius*. In my opinion, these species are much more similar in their morphology to the genus *Isohypsibius*, especially to the *Isohypsibius prosostomus* species complex. Considering these differences and in order to maintain morphological unity of the family Acutuncidae, I propose to move both species to the genus *Isohypsibius* (Isohypsibioidea, Isohypsibiidae) as *Isohypsibius parvus* (Lisi, Sabella & Pilato, 2014) **comb. nov.** and *Isohypsibius tibetanus* (Li & Li, 2008) **comb. nov.**

The diagnosis of the family Acutuncidae should be amended in order to include the genus *Mixibius* and formulated as follows: Hypsibioidea; pharynx with two macroplacoids, without microplacoids or a septulum; external (posterior) claws of the *Isohypsibius* type, internal (anterior) claws of the *Hypsibius* type (Pilato & Binda 2010).

The amended diagnosis of genus *Acutuncus* should be formulated as follows: Acutuncidae; AISMs in the form of hooks, asymmetrical with respect to the frontal plane, with the dorsal hook being shorter and higher than the ventral; caudal ends of AISMs usually blunt or rounded, not forming obviously sharp points; dorsal and ventral thickenings of the buccal tube posterior to the AISMs present; eggs with pillars in the chorion, visible under LM.

No distinctive morphological diagnosis can be given for the genus *Mixibius*. Apart from the molecular clade revealed in the present investigation, the genus *Mixibius* formally includes all species with the claws of the acutuncid type and without information on egg morphology and genetic data. Considering its diverse morphology (presence or absence of gibbosities of the body surface, different shape of AISMs), this taxon seems to be artificial and should be treated as tentative, waiting for revision.

### Notes on the phylogeny of the genus *Hypsibius*

The phylogenetic analysis of the superfamily Hypsibioidea (Fig. 9) revealed the presence of a fully supported clade for the family Hypsibiidae Pilato, 1969 (sensu Tumanov & Tsvetkova 2023), which incorporates all currently investigated genera of Diphasconinae Dastych, 1992 (*Diphascon* Plate, 1888, *Kararehius* Zawierucha, Stec & Shain, 2023 (in Zawierucha et al. 2023), and *Kopakaius* Zawierucha, Stec & Shain, 2023 (in Zawierucha et al. 2023)) and Hypsibiinae Pilato, 1969 (*Borealibius*, *Cryobiotus*, *Hypsibius*, and *Parahypsibius* Gąsiorek, 2024 in Gąsiorek *et al*. 2024). Relationships of the subclades within this monophyletic clade are not fully resolved (Fig. 10). There are two well-supported subclades: a clade comprising all genera of Diphasconinae and a clade comprising all genera of Hypsibiinae except for the genus *Parahypsibius*. The latter genus had an unstable position within the hypsibiid clade. In the Bayesian analysis, it formed a well-supported (posterior probability value of 0.99) monophyletic group with the diphasconin subclade, while in the ML analysis it is grouped with moderate support (bootstrap value of 87) with the Hypsibiinae subclade. This is in contrast to the results of Gąsiorek *et al*. (2024), where *Parahypsibius* was grouped with Hypsibiinae in both Bayesian and ML analyses. These differences can be explained by the different sets of taxa, selected for the analyses, by the different sets of analysed genes, or by the change of the outgroup to macrobiotoideans.

**FIGURE 10.**
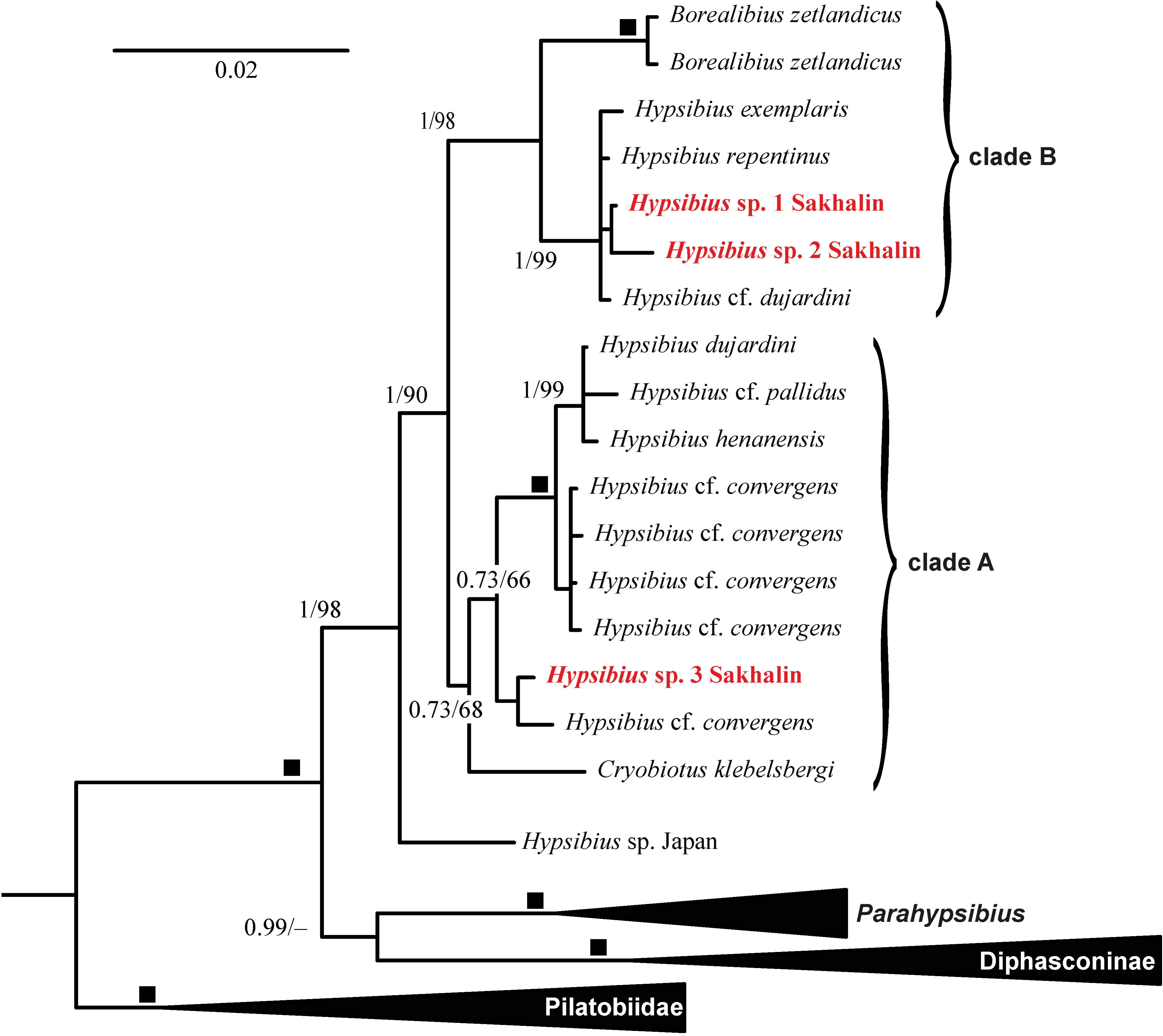
The phylogeny of Hypsibiidae based on concatenated 18S + 28S rRNA sequences. Numbers at nodes indicate Bayesian posterior probability values (BI, on the left) and bootstrap values (ML, on the right). Black squares indicate the nodes supported by values of 1/100% with both methods. Scale bar and branch lengths refer to the Bayesian analysis.

Within the Hypsibiinae clade, a single undescribed *Hypsibius* species from Japan (Ono *et al*. 2022) forms a sister lineage to all other species, which are distributed between two clades. The first (Fig. 10, clade A) comprises the genus *Cryobiotus* together with some species of the genus *Hypsibius*: *H. dujardini* (Doyère, 1840), *H. henanensis* Wang et al., 2024 (in Li *et al*. 2024), *H.* cf. *pallidus*, and several undescribed species of *H. convergens* morphogroup including *Hypsibius* sp. 3 from Sakhalin. Subcladal relations within this clade are not resolved. The second (Fig. 10, clade B) comprises the genus *Borealibius* together with another set of species of the genus *Hypsibius*: *H. exemplaris* Gąsiorek, Stec, Morek & Michalczyk, 2018a, *H. repentinus* Tumanov & Avdeeva, 2021, *H.* cf. *dujardini*, and two other *Hypsibius* species from Sakhalin (i.e., *Hypsibius* sp. 1 and *Hypsibius* sp. 2). Clade B is well-supported in my analysis, while clade A received poor support (0.73 in Bayesian analysis and 68 in ML), possibly because of lack of 28S rRNA data for most of the specimens, labelled as *H.* cf. *convergens*.

It is worth mentioning that the uncorrected genetic *p*-distances for the ITS-2 marker are noticeably low between the species within both clades A and B (not exceeding 3.2%), being much higher between the species belonging to the different clades (12–14%). This makes the ITS-2 marker useless for differentiating species within both clades. Comparison of the results of the bPTP species delimitation, performed on the basis of the COI and ITS-2 sequences confirmed this assumption (see SM.04 and SM.07).

These results are in concordance with results of the previous analyses (Tumanov and Tsvetkova 2023; Gąsiorek *et al*. 2024). The genus *Hypsibius* is evidently paraphyletic in relation to the genera *Cryobiotus* and *Borealibius*. It should be noted that both subclades in the *Hypsibius* clade *sensu lato* (including *Cryobiotus* and *Borealibius*) comprises species both with and without a septulum in the pharynx. Thus, it should be concluded that the presence or absence of this structure has low phylogenetic significance, so the terms *Hypsibius dujardini* group and *Hypsibius convergens* group (the only difference between these groups is the presence or absence of the septulum) can be used only as designations of morphogroups within the genus (Gąsiorek *et al*. 2024).

### Conclusion

Despite the small number of samples and specimens studied, the results obtained during this study are of certain faunistic and phylogenetic interest. The number of species recorded for Sakhalin Island increased from 1 to 9, and what is much more significant – nearly all species found seem to be new to science. Their formal description is not possible because of the small number of specimens obtained. This is evidence of the substantial distinctness of the Far East fauna of freshwater tardigrades from the relatively well-studied European fauna. Molecular identification revealed the presence of potentially cryptic diversity within the *Hypsibius dujardini* morphogroup in the Sakhalin fauna. Inclusion of the obtained molecular data in the phylogenetic analysis of the superfamily Hypsibioidea revealed presence of the well-supported clade of the enigmatic genus *Mixibius* for the first time, and made it possible to recognize its phylogenetic position as the sister clade to the genus *Acutuncus* within the family Acutuncidae.

## Supplementary materials

SM.01 – Complete list of sequences used in phylogenetic analysis.

SM.02 – Alignment of the concatenated 18S rRNA and 28S rRNA sequences used for the phylogenetic analysis.

SM.03 – Uncorrected genetic *p*-distances for COI and ITS-2 sequences between the studied specimen and sequences of the genus *Hypsibius* (with addition of the closely related genera *Borealibius* and *Cryobiotus*) available in GenBank.

SM.04 – Species delimitation tree for the genus *Hypsibius* generated by the Bayesian Poisson Tree Processes (bPTP) model, using a fragment of the COI gene. Black lines indicate branching processes among species, red lines indicate branching processes within species.

SM.05 – Uncorrected genetic *p*-distances for COI and ITS-2 sequences between the studied specimen and sequences of the genus *Mesobiotus* available in GenBank.

SM.06 – Species delimitation tree for the genus *Mesobiotus* generated by the Bayesian Poisson Tree Processes (bPTP) model, using a fragment of the COI gene. Black lines indicate branching processes among species, red lines indicate branching processes within species.

SM.07 – Species delimitation tree for the genus *Hypsibius* generated by the Bayesian Poisson Tree Processes (bPTP) model, using ITS-2 marker. Black lines indicate branching processes among species, red lines indicate branching processes within species.

## Acknowledgements

The expedition to Sakhalin and the field work was supported by the non-profit charitable foundation “Support of bioresearch “BIOM”, grant № 4/2024-гр. The laboratory work conducted in Saint Petersburg State University was supported by the Russian Science Foundation, grant No. 25-74-20033 “Evolutionary transformations of nanostructural elements of the Ecdysozoa cuticle using the example of the integumentary structures of tardigrades (Tardigrada)”. I am grateful to Olga Knyazeva for the linguistic review of the manuscript. I am thankful to anonymous reviewers for the valuable comments and corrections. The laboratory work was conducted using the equipment of the Core Facilities Centre “Centre for Molecular and Cell Technologies” of Saint Petersburg State University (https://researchpark.spbu.ru/index.php/en/biomed-eng) and “Taxon” Research Resource Center of the Zoological Institute of the Russian Academy of Sciences (http://www.ckp-rf.ru/ckp/3038/).

## Notes

### Competing Interest Statement

The authors have declared no competing interest.

