## Supplementary material for "Set to its place: first DNA data on freshwater tardigrades of Sakhalin (Far East Russia) shed light on the phylogenetic position of *Mixibius* (Eutardigrada: Hypsibioidea)": SM.01

Table 1. Complete list of sequences used in the phylogenetic analysis. Sequences produced in this study are marked in bold.

| Species | 18S rRNA | 28S rRNA | Source |
| --- | --- | --- | --- |
| Hypsibioidea |  |  |  |
| Ramazzottiidae |  |  |  |
| *Ramazzottius oberhaeuseri* | MG573241 | MG573242 | Stec et al. 2018b |
| *Ramazzottius varieornatus* | AP013352 | MG432818 | Hashimoto et al. 2016, Zawierucha et al. 2018 |
| *Cryoconicus kaczmareki* | MG432796 | MG432797 | Zawierucha et al. 2018 |
| *Hebesuncus conjungens* | PQ108465 | PQ108478 | Vecchi & Stec 2025 |
| *Hebesuncus ryani* | EU266956 |  | Sands et al. 2008 |
| Calohypsibiidae |  |  |  |
| *Calohypsibius ornatus* | OM304865 | OM304872 | Tumanov & Tsvetkova 2023 |
| Microhypsibiidae |  |  |  |
| *Microhypsibius* sp. 1 | OM304866 | OM304873 | Tumanov & Tsvetkova 2023 |
| *Microhypsibius* sp. 2 | OM304867 | OM304874 | Tumanov & Tsvetkova 2023 |
| Acutuncidae |  |  |  |
| *Acutuncus antarcticus* | OM278641 | OM278644 | Vecchi et al. 2023 |
| *Acutuncus giovanniniae* | OM350035 | OM350031 | Vecchi et al. 2023 |
| *Acutuncus mariae* | MW012739 | MW012742 | Zawierucha et al. 2020 |
| *Acutuncus mecnuffi* | OM350033 | OM350029 | Vecchi et al. 2023 |
| *Mixibius saracenus* | HQ604955 |  | Bertolani et al. 2014 |
| *Mixibius* cf. *saracenus* | HQ604956 |  | Bertolani et al. 2014 |
| ***Mixibius* sp.** | **PX243403** | **PX232532** | **this study** |
| Pilatobiidae |  |  |  |
| *Fontourion recamieri* | KX347526 | KX347527 | Gąsiorek et al. 2017 |
| *Fontourion islandicum* | MH682258 | MH682257 | Buda et al. 2018 |
| *Fontourion glacialie* | MW012733 | MW012743, MW012744 | Zawierucha et al. 2020 |
| *Pilatobius bullatus* | OM304862 | OM304869 | Vecchi et al. 2023 |
| *Degmion nuominense* | MT330115 | MT330116 | Sun et al. 2021 |
| *Notahypsibius pallidoides* | MN912103 | MK967962 | Tumanov 2020 |
| Itaquasconidae |  |  |  |
| *Guidettion prorsirostre* | MT126750–1 | MT126768–9 | Gąsiorek & Michalczyk 2020 |
| *Guidettion* sp. | OM304861, OQ351310 | OM304868, OQ357539 | Vecchi et al. 2023, Tumanov & Tsvetkova 2023 |
| *Adropion scoticum* | MT126752–3, OP035715 | MT126770–1, OP035794 | Gąsiorek & Michalczyk 2020, Tumanov et al. 2022 |
| *Adropion belgicae* | HQ604925–6 |  | Bertolani et al. 2014 |
| *Adropion sp n 1_PL 276* | MT126748 | MT126766 | Gąsiorek & Michalczyk 2020 |
| *Adropion sp n 2_NO 018* | MT126749 | MT126767 | Gąsiorek & Michalczyk 2020 |
| *Arctodiphascon tenue* | OQ351311–16 | OQ357540–45 | Tumanov & Tsvetkova 2023 |
| *Mesocrista spitzbergensis* | KX347532 | KX347533 | Gąsiorek et al. 2016 |
| *Mesocrista revelata* | KU528627, OP035717 | KX347536, KU528630–1, KU528628, OP035797 | Gąsiorek et al. 2016, Tumanov et al. 2022 |
| *Platicrista* sp. | OQ351317 | OQ357546 | Tumanov & Tsvetkova 2023 |
| *Platicrista* aff*. angustata* | MT126760–2 | MT126779–81 | Gąsiorek & Michalczyk 2020 |
| *Platicrista horribilis* | MT126763 | MT126782 | Gąsiorek & Michalczyk 2020 |
| *Astatumen trinacriae* | FJ435731–3 | FJ435773–5 | Guil & Giribet 2012 |
| *Astatumen bartosi* | MT126754 | MT126772 | Gąsiorek & Michalczyk 2020 |
| *Astatumen* sp. | OQ351318 | OQ357547 | Tumanov & Tsvetkova 2023 |
| *Astatumen* sp. | OQ351319 | OQ357548 | Tumanov & Tsvetkova 2023 |
| *Raribius minutissimus* | MT126758 | MT126777 | Gąsiorek & Michalczyk 2020 |
| *Insulobius orientalis* | MT126764 | MT126783 | Gąsiorek & Michalczyk 2020 |
| *Itaquascon serratulum* | MT126759 | MT126778 | Gąsiorek & Michalczyk 2020 |
| Hypsibiidae |  |  |  |
| *Borealibius zetlandicus* | HQ60492–4 |  | Bertolani et al. 2014 |
| *Hypsibius convergens* | FJ435726 | FJ435771 | Guil & Giribet 2012 |
| *Hypsibius dujardini* | MG777532 | MG777533 | Gąsiorek et al. 2018 |
| *Hypsibius exemplaris* | HQ604943 | MG800337 | Bertolani et al. 2014, Gąsiorek et al. 2018 |
| *Hypsibius henanensis* | OR557238 | OR557239 | Li et al. (2024) |
| *Hypsibius pallidus* | HQ604945 |  | Bertolani et al. 2014 |
| *Hypsibius repentinus* | MN927184 | MW549064 | Tumanov & Avdeeva 2021 |
| *Hypsibius* cf. *convergens* | KC582832, PP736631–3 |  | Dabert et al. 2014, Zhang et al. 2024 |
| *Hypsibius* cf. *dujardini* | PP736635 |  | Zhang et al. 2024 |
| *Hypsibius* sp. Japan | ON898549 | ON927924–5 | Ono et al. 2022 |
| ***Hypsibius* sp. 1** | **PX243404** | **PX232533** | **this study** |
| ***Hypsibius* sp. 2** | **PX243405** | **PX232534–5** | **this study** |
| ***Hypsibius* sp. 3** | **PX243406** | **PX232536** | **this study** |
| *Cryobiotus klebelsbergi* | KT901827 | KC582835 | Dabert et al. 2014, 2015 |
| *Parahypsibius scabropygus* | AM500649, KC582831, PQ421023 | PQ421035 | Kiehl et al. 2007, Dabert et al. 2014, Gąsiorek et al. 2024 |
| *Parahypsibius* sp. | PQ421031 | PQ421043 | Gąsiorek et al. 2024 |
| *Diphascon pingue* | FJ435734, FJ435736 | FJ435776–7 | Guil & Giribet 2012, |
| *Diphascon* cf*. pingue* | OP035716, OQ351320–1 | OP035795–6, OQ357549 | Tumanov et al. 2022, Tumanov & Tsvetkova 2023 |
| *Diphascon higginsi* | HQ604932 |  | Bertolani et al. 2014 |
| *Diphascon mauccii* | EU266945 |  | Sands et al. 2008 |
| *Diphascon puniceum* | EU266946–9 |  | Sands et al. 2008 |
| *Diphascon wuyingensis* | MK387067 |  |  |
| *Diphascon greveni* | OQ351322–6 | OQ357550–4 | Tumanov & Tsvetkova 2023 |
| *Kopakaius nicolae* | OP191650 |  | Zawierucha et al. 2023 |
| *Kopakaius* sp. 1 | OP191642, OP191646 | OP191651, OP191654 | Zawierucha et al. 2023 |
| *Kararehius gregorii* | OP191647–8 | OP191640–1 | Zawierucha et al. 2023 |
| Macrobiotoidea |  |  |  |
| *Richtersius coronifer* | MH681760 | MH681757 | Stec et al. 2020 |
| *Macrobiotus shonaicus* | MG757132 | MG757133 | Stec et al. 2018a |

<https://doi.org/10.11646/zootaxa.5492.1.5>
