## Supplementary figures and images for "Set to its place: first DNA data on freshwater tardigrades of Sakhalin (Far East Russia) shed light on the phylogenetic position of *Mixibius* (Eutardigrada: Hypsibioidea)"

### SM.04

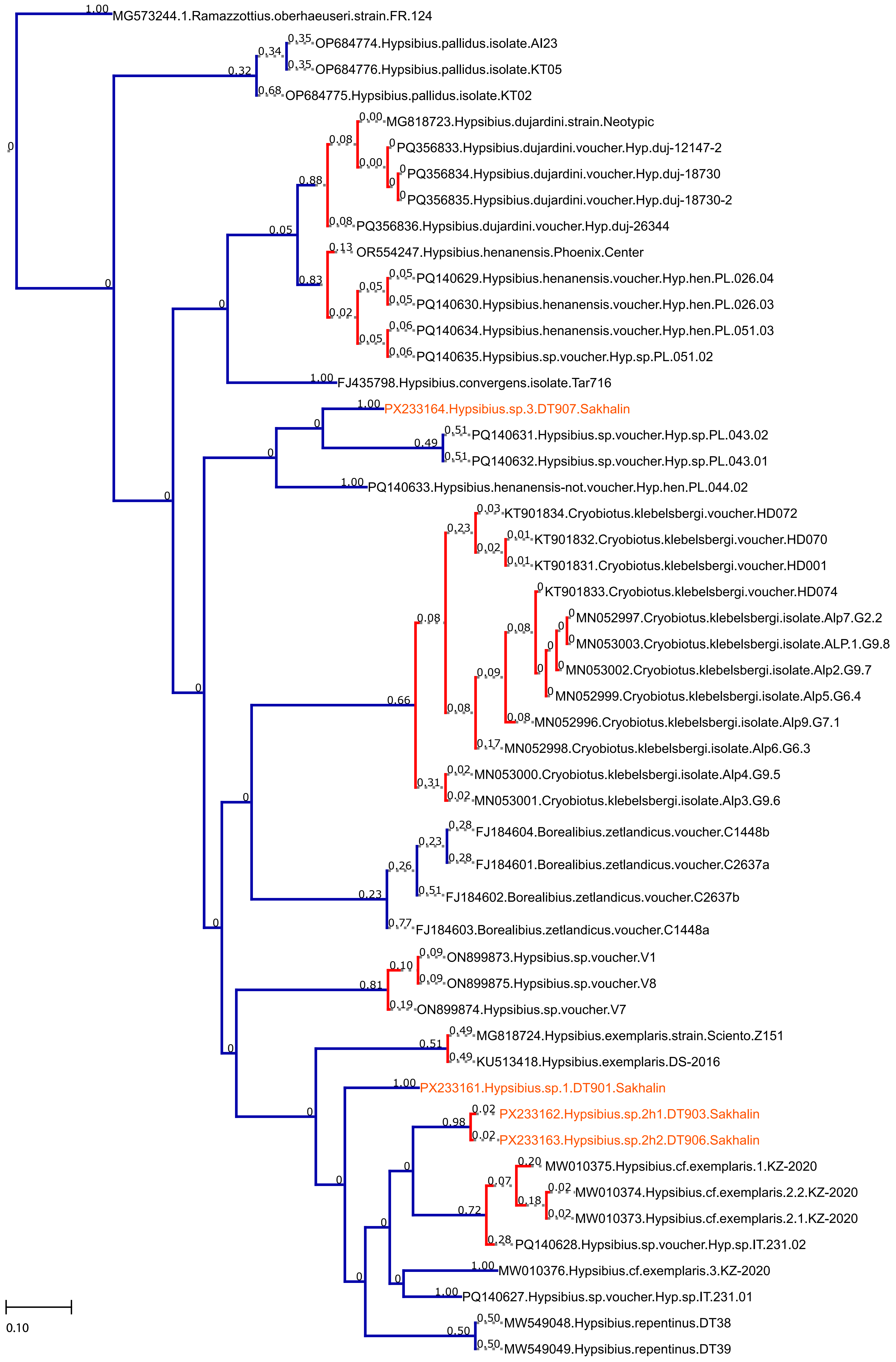

### SM.06

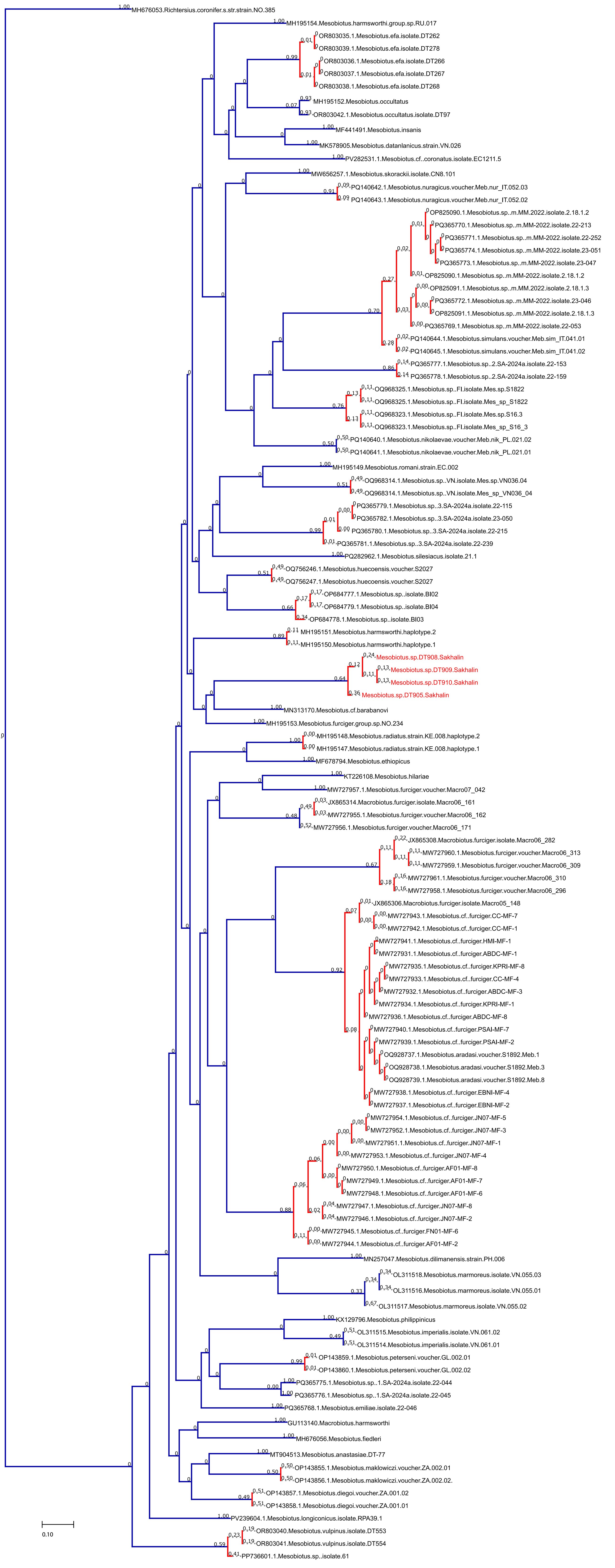

### SM.07

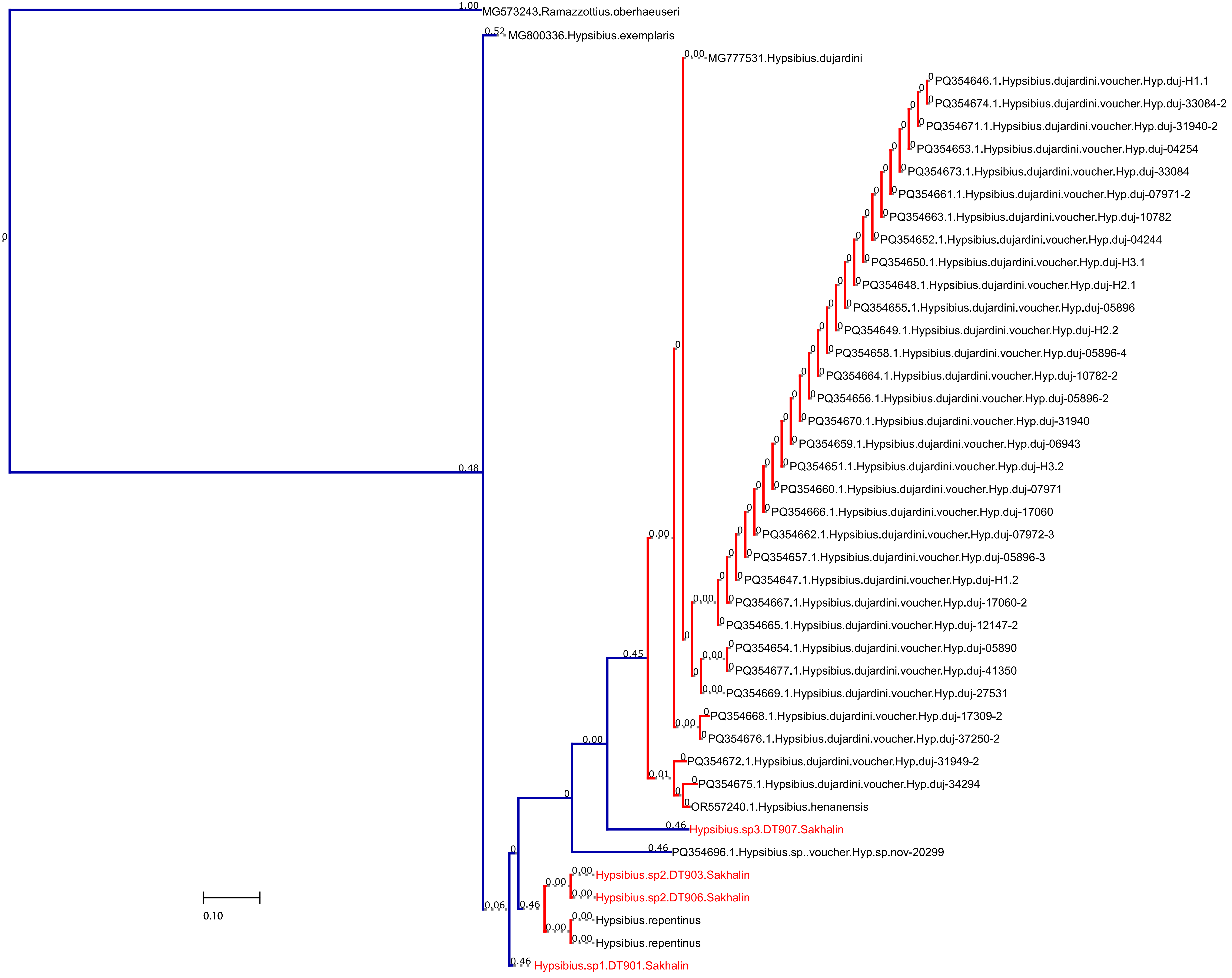
